# Entrainment of local synchrony reveals a causal role for high-beta right frontal oscillations in human visual consciousness

**DOI:** 10.1101/574939

**Authors:** Marine Vernet, Chloé Stengel, Romain Quentin, Julià L. Amengual, Antoni Valero-Cabré

**Affiliations:** Institut du Cerveau et de la Moelle Epinière (ICM), CNRS UMR 7225, INSERM U 1127 and Université Pierre et Marie Curie, Paris, France; Laboratory for Cerebral Dynamics Plasticity and Rehabilitation, Boston University, School of Medicine, Boston, MA, USA; Cognitive Neuroscience and Information Technology Research Program, Open University of Catalonia (UOC), Barcelona, Spain

## Abstract

Prior evidence supports the critical role of oscillatory activity in cognitive function, but are cerebral oscillations simply correlated or causally linked to specific aspects of visual cognition? Here, EEG signals were recorded on humans performing a conscious visual detection task, while they received brief *rhythmic* or *random* noninvasive stimulation patterns delivered to the right Frontal Eye Field prior to the onset of a lateralized target. Compared to *random* patterns, *rhythmic* high-beta patterns led to greater entrainment of local oscillations (i.e., increased power and phase alignment at the stimulation frequency), and to higher conscious detection of contralateral targets. When stimulation succeeded in enhancing visual detection, the magnitude of oscillation entrainment correlated with visual performance increases. Our study demonstrates a causal link between high-beta oscillatory activity in the Frontal Eye Field and conscious visual perception. Furthermore, it supports future applications of brain stimulation to manipulate local synchrony and improve or restore impaired visual behaviors.

## Introduction

Within the last decade, there has been increasing interest in the involvement of oscillations and synchronization in information coding (Buzsaki and Draguhn, 2004; Fries et al., 2007). In the research field of attention and perception, occipital or parietal alpha (Thut et al., 2006; Marshall et al., 2015) and theta (Dugue and VanRullen, 2017), fronto-parietal beta (Buschman and Miller, 2007; Phillips and Takeda, 2009), occipital and fronto-parietal gamma synchrony (Buschman and Miller, 2007; Wyart and Tallon-Baudry, 2009; Marshall et al., 2015) and fronto-parietal rhythmic states coordinated by theta oscillations (Fiebelkorn et al., 2018) have been associated with the ability to acknowledge the presence of a visual target.

Additionally, local and interregional oscillations at these frequency bands are thought to code for specific cognitive processes. For example, in monkeys, episodes of fronto-parietal synchronization at 30 and 50 Hz have been correlated with top-down and bottom-up attention orientation, during a visual search task and a pop-out visual task, respectively (Buschman and Miller, 2007). Similarly in humans, 30-Hz rhythmic patterns of Transcranial Magnetic Stimulation (TMS) delivered to a right frontal region increased conscious visual sensitivity, suggesting that high-beta synchrony (Chanes et al., 2013) within a fronto-parietal network (Quentin et al., 2015; Quentin et al., 2016) is causally involved in conscious visual perception. However, are these behavioral effects causally mediated by the engagement of frequency-specific oscillatory activity? Or, do cortical oscillations simply represent an epiphenomenon devoid of any direct implication concerning visual cognition and associated visually-guided behaviors (Pareti and De Palma, 2004)?

Two noninvasive brain stimulation methods, rhythmic transcranial magnetic stimulation (rhythmic TMS) (Thut and Pascual-Leone, 2010; Thut et al., 2011a) and transcranial alternating current stimulation (tACS) (Antal and Paulus, 2013), have been used to manipulate oscillatory activity in humans. While tACS is portable, easy to use and well suited to synchronize wide cortical regions in view of clinical applications, rhythmic TMS capitalizes on its higher focality and unique ability to induce brief oscillatory effects (in trial-by-trial designs). This enables testing of relevant causal contributions of rhythmic activity at specific time-windows during human cognitive processing.

To address the crucial question of the causal role of local oscillations in conscious perception, we designed an experiment in which participants performed a conscious visual task while receiving rhythmic TMS patterns. Such patterns were expected to entrain short episodes of high-beta rhythmic activity (30 Hz) in the Frontal Eye Field (FEF), a key node of the dorsal attentional network (Corbetta et al., 2008) involved in the modulation of visual perception in monkeys (Moore and Fallah, 2001) and humans (Grosbras and Paus, 2002, 2003; Chanes et al., 2012). Combined TMS-EEG recordings have shown the entrainment of short-lasting alpha oscillations (∼10 Hz) by noninvasive rhythmic stimulation delivered at alpha frequency to posterior parietal locations at rest, i.e., in participants not engaged in any specific task-driven behavior (Thut et al., 2011b). Beta frequencies in motor or memory systems have been successfully entrained in recent studies (Hanslmayr et al., 2014; Romei et al., 2016b). However, entrainment at such frequencies over regions specifically relevant for visual cognition and conscious access remains to be explored.

In the present study, a group of right-handed healthy human participants performed a conscious visual detection task. Before the onset of the visual target, either active or sham TMS bursts were applied over the right FEF. We compared the neurophysiological (*EEG recordings*) and behavioral (*conscious visual detection performance*) effects of *rhythmic* stimulation patterns, composed of 4 TMS pulses at 30 Hz, to the effects of *random* TMS patterns (with identical time onset, duration, and pulse number, but providing arrhythmic activity). We hypothesized that *rhythmic*, but not *random*, right frontal TMS patterns would progressively phase-align local oscillatory activity at the frequency band dictated by the temporal organization of the input stimulation patterns (Thut et al., 2011b; Thut et al., 2017). More importantly, we predicted improvements of visual performance under the impact of high-beta rhythmic patterns, and eventually an association between the magnitude of evoked high-beta oscillations and conscious visual detection improvements. These results would provide evidence in favor of a causal contribution of oscillatory activity to specific aspects of visual cognition. It would also support future manipulations of local synchrony to improve or restore specific aspects of perceptual performance, subtended by oscillation-based coding mechanisms.

## Materials and Methods

### Participants

Fourteen right-handed participants (9 women and 5 men) between 20 and 34 years old (24±4) took part in the study. Participants underwent all experimental conditions (within-subject experimental design), were naïve to the purpose of the experiment, reported no history of neurological or psychiatric disorders, had normal or corrected-to-normal vision, and participated voluntarily after providing written informed consent. The protocol (C08-45) was reviewed by the Inserm (Institut National de la Santé et la Recherche Scientifique) ethical committee and was approved by an Institutional Review Board (CPP Ile de France 1) in accordance with the Declaration of Helsinki.

### Apparatus

Participants were seated with their head positioned on a chin-rest and their eyes directed towards a computer screen (22’’) 57 cm away. A custom-made script, using the MATLAB (Mathworks) Psychtoolbox (Brainard, 1997), ran on a desktop computer (HP Z800, Hewlett Packard). It synchronized the presentation of visual stimuli on the computer screen, the pulses delivered by two biphasic repetitive TMS devices (SuperRapid, Magstim) attached to standard 70 mm figure-of-eight coils operated via a trigger-synchronization device (Master 8, A.M.P.I.), a remote gaze tracking capture system (Eyelink 1000, SR Research), and EEG recordings performed with TMS-compatible equipment (BrainAmp DC, BrainVision Recording Software, EasyCap and Ag/AgCl sintered ring electrodes, BrainProducts GmbH). Additionally, a frameless neuronavigation system (Brainsight, Rogue Research) was used throughout the experiment to deliver TMS on precise standardized coordinates corresponding to the right FEF.

### Conscious visual detection paradigm

A visual detection paradigm similar to the one employed in prior studies was used (Chanes et al., 2012; Chanes et al., 2013; Chanes et al., 2015; Quentin et al., 2015; Quentin et al., 2016). Each session included a titration block, a training block, and 4 experimental blocks. The latter blocks assessed the effects of rhythmic/random TMS on EEG signals and visual detection (2 blocks for each stimulation pattern). Each block was divided into sub-blocks of 20 trials. Calibration and training blocks were tailored in length to each participant and completed once stable performance was reached. Experimental blocks included a fixed number of 140 trials divided in 7 sub-blocks.

Each trial (see Figure 1A) started with a gray resting screen (luminance: 31 cd/m2, 2500 ms) followed by a fixation screen (randomly lasting between 1000 and 1500 ms). A fixation cross (0.5×0.5°) was displayed at the center of the screen along with two lateral placeholders (6.0×5.5°, eccentricity 8.5°). The fixation cross became slightly larger (0.7×0.7°, 66 ms) to alert participants of an upcoming event. Then, following a fixed inter-stimulus interval (233 ms), a target could appear for a brief period of time (33 ms) in the center of one of the two placeholders (40% left trials, 40% right trials, 20% no target or “catch” trials). The target was a low-contrast Gabor stimulus made of vertical lines (0.5°/cycle sinusoidal spatial frequency, 0.6° exponential standard deviation, minimum and maximum Michelson contrast of 0.005 and 1, respectively).

**Figure 1.**
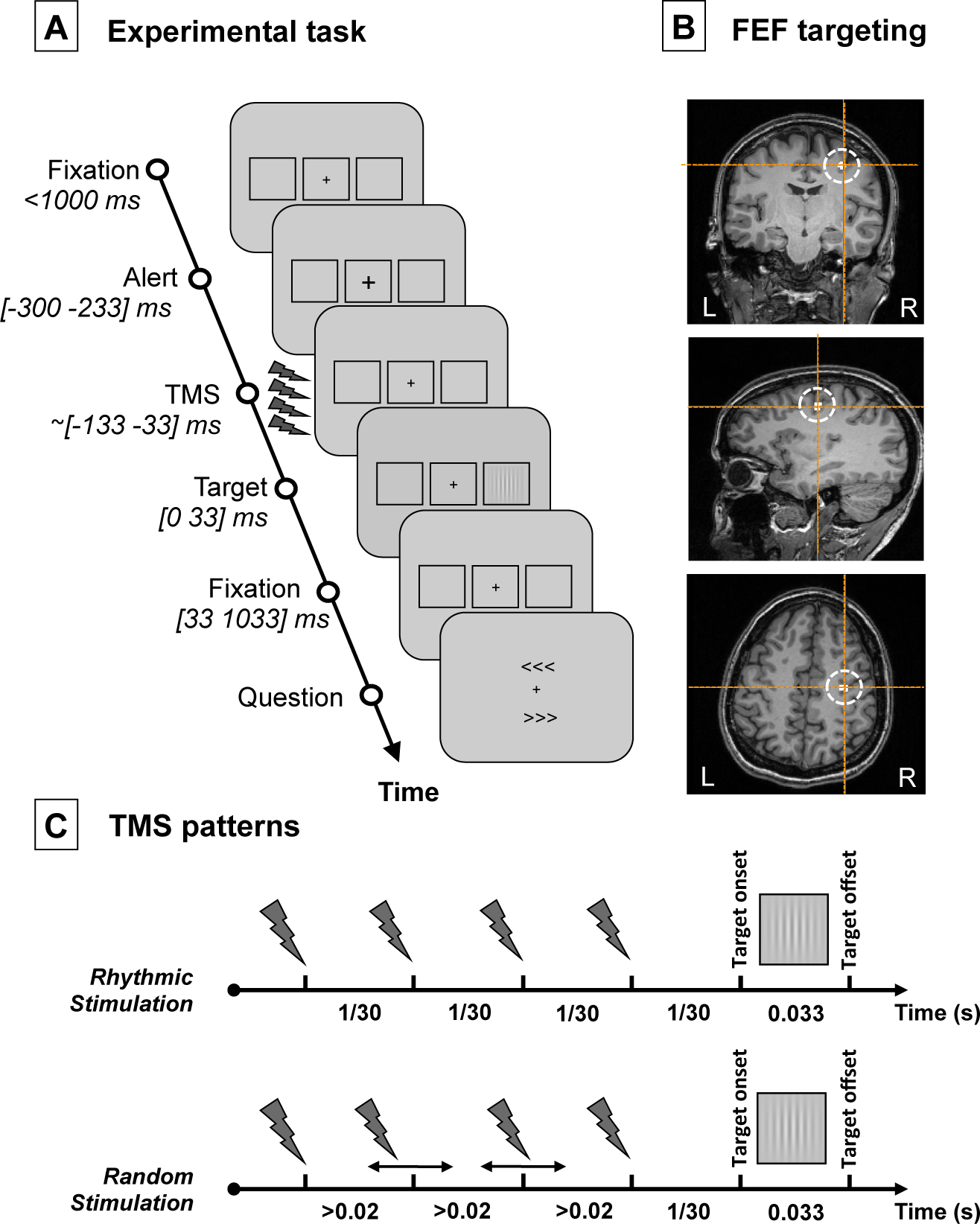
Experimental task, TMS targeted cortical region and stimulation patterns. **(A)** Visual detection task performed by participants. After a period of fixation, a central cross became slightly larger to alert participants of an upcoming event. Then active/sham rhythmic/random TMS patterns were delivered to the right FEF region prior to the presentation of a visual target at the center of a right/left placeholder. Participants were requested to indicate whether they did or did not perceive a target and, if they did, where it appeared (no target perceived/target perceived on the right/target perceived on the left). Notice that in 20% of the trials (“catch trials”), no target was presented in any of the two placeholders; **(B)** Coronal, sagittal and axial T1-3D MRI sections from a representative participant generated by the frameless stereotaxic neuronavigation system showing the localization of the right FEF, stimulated in our experiment (Talairach coordinates X=31, Y=-2, Z=47 (Paus, 1996)); **(C)** Schematic representation of the temporal distribution of the 4-pulse bursts employed for the 30 Hz *rhythmic* and the *random* stimulation conditions. Contrasting the behavioral (conscious visual performance) and electrophysiological (EEG) impact elicited by these two patterns isolates the effects of 30 Hz FEF activity (only present in rhythmic bursts) from those induced by 4 TMS pulses delivered during a 100 ms interval (featured by both rhythmic and random bursts).

As in prior studies (Chanes et al., 2012; Chanes et al., 2013; Quentin et al., 2015), participants were asked to report whether they consciously perceived the target or not and, if they did, where in the screen it appeared (detection task). To do so, two arrow-like signs (“<<<” and “>>>”), pointing to the left and to the right, were simultaneously presented below and above the fixation cross. Participants used 3 keys of a keyboard to answer: an upper key “d”, a lower key “c” and the space bar, which they operated with the middle, index, and thumb fingers of their left hand, respectively. Participants were requested to respond either by pressing the space bar if they did not see the stimulus, or by pressing “d”/“c” to select the upper/lower arrow-like sign pointing to the placeholder where they had perceived the target. The location of each arrow, above or below the fixation point, was randomized across trials. The response of the participant ended the trial.

Based on previously used procedures (Cornsweet, 1962; Chanes et al., 2013), a titration block served to estimate the contrast for each participant where ∼50% of the visual targets were consciously detected and reported (visual detection threshold). This procedure was performed under the effect of sham TMS patterns delivered before the target onset to the right FEF, identical to those used during the experiment (see full details on the TMS procedure section below). Participants initiated the titration with a high contrast target. A one-up/one-down staircase procedure was employed to adjust stimulus contrast in search of the threshold. The initial contrast step was equal to the initial contrast level. Then, contrast steps were divided by two on each reversal, but were always kept higher than 0.005 Michelson contrast units. We considered the 50% threshold to be reached when the last five consecutively tested contrasts were not different by more than 0.01 Michelson contrast units. This procedure was repeated twice. If the difference between the two estimated thresholds was lower than 0.01 Michelson units, the calibration procedure was terminated at the end of the ongoing sub-block and the average of the two last thresholds was taken as the final 50% threshold. If the difference between the two measures proved equal to or higher than 0.01 Michelson contrast units, the threshold was determined again. A short break was allowed at the end of each sub-block of testing.

During the ensuing training block, active or sham TMS was delivered on the right FEF (see further details on the TMS procedure section below). Half of the trials for each condition (left target, right target, no target) were performed with active TMS, whereas the other half were performed with sham TMS. Within each block, all types of trials, i.e., three visual targets (left, right, no target) performed under two TMS conditions (real, sham), were carried out in randomized order. This training block aimed to further familiarize the participant with the stimulation and check the consistency of visual detection performance of sham trials (previously titrated at 50% correct detections under sham TMS) when both active and sham TMS trials were randomly mixed-up during the same block. At the end of each sub-block, participants were allowed to take a rest and the experimenter re-adjusted target contrast if necessary.

Once participants carried out the training task consistently and according to the established titration level, they were invited to perform 4 experimental blocks of 140 trials (each organized in 7 sub-blocks of 20 trials). These were identical to the training block, except that the target contrast was kept constant and short breaks were allowed every two sub-blocks.

For fixation control purposes, eye movements were monitored. Fixation was considered broken when participants’ gaze was recorded outside a 2° radius of the fixation cross any time between cue onset and target offset. When this occurred, participants received an alert message; that specific trial was randomized with the rest of the trials left in the sub-block and repeated. At each break, participants received an alert signal if their false alarm rate (i.e., reporting having seen a target when no target was presented) was higher than 50% (only during the calibration and training blocks), and they were informed on the percentage of target location errors and on the percentage of incorrect gaze central fixations (at the end of each block and throughout the experiment).

### Noninvasive brain stimulation procedures with rhythmic TMS

Active TMS bursts were delivered to the right FEF. The right FEF region (see Figure 1B) was localized on each individual T1-weighted MRI scan (3T Siemens MPRAGE, flip angle=9, TR=2300 ms, TE=4.18 ms, slice thickness=1mm, isovoxel) using averaged Talairach coordinates x=31, y=-2, z=47 (Paus, 1996) and a 0.5 cm radius spherical Region of Interest (ROI). At all times, the TMS coil was held tangentially to the skull, with its handle oriented caudal-to-rostral and medial-to-lateral at ∼45° with respect to the longitudinal interhemispheric fissure (i.e., ∼parallel to the central sulcus), and kept within an area of ∼1-2 mm radius from the right FEF by means of a frameless neuronavigation system.

Sham TMS bursts were randomly interleaved during the same block. They were delivered by a second TMS coil, placed next to the right FEF site and oriented perpendicular to the scalp, which prevented the magnetic field from reaching the skull and stimulating the brain. Acoustic stimulation related to the coil discharge noise was diminished by having the participant wearing earplugs; skull bone vibration when the coil is discharging can also contribute to auditory evoked potentials and entrainment (for a review, see (Vernet and Thut, 2014), see also (ter Braack et al., 2015)). Nevertheless, those two concerns were limited by having the sham coil also in contact with the skull, thus mimicking the accompanying auditory and somatosensory effects of active TMS, even if some slight differences remain (Thut et al., 2005). Furthermore, to minimize attentiveness to the TMS itself, participants were familiarized during the training with the sensations associated to transcranial stimulation. It should be also noted that during the experiments itself, participants were largely attentive to the challenging perceptual task to be performed.

Stimulation consisted of either rhythmic or random bursts made of 4 consecutive TMS pulses with a total duration of 100 ms. The last pulse was delivered 1/30 second (i.e., a cycle of a 30 Hz oscillation) prior to the onset of the visual target. Rhythmic patterns consisted of four pulses uniformly distributed to produce a regular 30 Hz burst. Random patterns, which were used as control patterns to isolate the specific contribution of stimulation frequency, had their 1^st^ and 4^th^ pulses delivered at the same timing as in the rhythmic patterns. In contrast, the 2^nd^ and 3^rd^ pulses of the burst were randomly delivered in the interval left between the 1^st^ and 4^th^ pulses respecting the following limitations: (1) a minimum interval of 20 milliseconds between two contiguous pulses, to ensure the effective recharge of TMS machines capacitors; and (2) a minimum interval of three milliseconds between the timing of each of the two middle pulses and the timing that would have been held in pure 30 Hz rhythmic patterns (see Figure 1C). Rhythmic and random TMS were delivered in separated experimental blocks performed in a counterbalanced order (7 participants started with rhythmic TMS block and 7 with random TMS block).

Our TMS design (within-block active/sham conditions; between-block rhythmic/random conditions) has been successfully tested in several TMS studies combining rhythmic stimulation and perceptual behaviors (Chanes et al., 2013; Chanes et al., 2015; Quentin et al., 2015; Quentin et al., 2016) in absence of EEG recordings. The type of stimulation delivered on each trial (*active* vs. *sham*) was randomized online by the computer in control of the behavioral paradigm. This within-block active/sham design controls for natural fluctuations of sustained attention and arousal level across the session.This provides an opportunity to subtract the potential impact caused by the unspecific effects associated to TMS delivery. This also precluded any possibility for neither participants nor the TMS operator to anticipate the type of TMS (*active* vs. *sham*) delivered on a given trial, hence protecting the experiment from conscious/unconscious biases. Interestingly, our preliminary testing revealed that when tested in separate blocks, differences between *rhythmic* vs. *random* TMS patterns passed unnoticed to participants. Indeed, debriefing performed in past reports (Chanes et al., 2013; Chanes et al., 2015; Quentin et al., 2015; Quentin et al., 2016) and also in the current study confirmed that participants (whose attention during the session was captured by a highly demanding near-threshold visual detection task) were unaware about the slight temporal differences of the delivered stimulation patterns (rhythmic vs. random) tested in separated blocks and presented in counterbalanced order.

Stimulation intensity was set up at a fixed value of 55% of the maximal stimulator output (MSO), instead of being adjusted to the individual resting motor threshold (RMT). Scalp-to-cortex distance is known to account for variability of the motor threshold (Stokes et al., 2005); nevertheless, other factors are probably at play for determining the excitability of other brain areas. Indeed, TMS-measured excitability in M1 poorly predicts the excitability of other areas (Stewart et al., 2001; Kahkonen et al., 2005). Our previous studies demonstrated behavioral effects at the group level using fixed intensity of 45% MSO (Chanes et al., 2013; Quentin et al., 2015). In the present study, the intensity of 55% took into account the estimated increased coil-to-cortex distance due to the presence of the EEG cap. However, to allow comparison with other studies (Rossi et al., 2009), the resting motor threshold (RMT) for the left *abductor pollicis brevis* (APB) muscle was determined on each participant at the end of the experiment as the minimum intensity at which TMS pulses applied on the right primary motor cortex (M1) yielded an activation of the APB in at least 50% of the attempts (RMT=72±9% MSO). The stimulation intensity applied to the right FEF and employed in the experiment corresponded on average to 78±12% of each participant’s motor threshold. Post-hoc analyses confirmed that the magnitude of the behavioral and physiological effects reported in the present study was not significantly correlated with individual RMT (p > 0.69).

### EEG recordings and analyses

EEG activity was continuously acquired from 60 scalp electrodes with the reference placed on the tip of the nose and the ground located on the left earlobe. Electrooculogram (EOG) was recorded with 4 additional electrodes (on the right and left temples and above and below one eye). Skin/electrode impedance was maintained below 5 kOhm. The signal was digitized at a sampling rate of 5000 Hz.

EEG signals were analyzed with MATLAB (R2013a), EEGLAB (v10.2.5.5.b) (Delorme and Makeig, 2004) and FieldTrip (Oostenveld et al., 2011) according to the following procedure. First, EEG data were epoched around the onset of the visual target ([-2 s, +2 s]). Then a 2^nd^ order infinite impulse response (IIR) Butterworth filter (pass-band between 1 and 50 Hz) was used with a forward-backward filtering to maintain a zero phase shift. This filter was not applied to time windows containing TMS pulses ([-4 ms, +12 ms] centered at each of the four pulses’ onset). Afterwards, TMS electromagnetic artifacts were eliminated from the EEG signals by removing this time window and performing a linear interpolation (Fuggetta et al., 2006; Vernet et al., 2014a; Vernet and Thut, 2014). The length of this interval (16 ms for each of the four pulses) was chosen after examination of the raw data in every participant. Data were then resampled at 500 Hz. Trials contaminated by blinks or muscle artifacts were identified by visual inspection, and removed. For each experimental condition, residual artifacts were segregated from physiological responses using a common Independent Component Analysis (ICA). For each participant, 8±4 components (range from 3 to 16) related to electrical artifacts were identified by activity strongly peaking in the vicinity of the stimulation site shortly after each TMS pulse, and by a spectrum covering a restricted frequency range with strong harmonics (Hamidi et al., 2010; Vernet et al., 2013). Once these components were removed, cleaned EEG signals were calculated back at the electrode level. This procedure of TMS artifacts removal was applied to all EEG trials, whether the magnetic stimulation applied was active or sham.

Once data cleaning was completed, two types of procedures were performed for each analysis: the first one included data from all 60 EEG scalp electrodes, whereas the second one concerned only EEG signals from the closest electrode to the stimulated right FEF area (i.e., electrode FC2).

The procedure to evaluate power (estimating the amount of rhythmic activity) and inter-trial coherence (evaluating the phase alignment of rhythmic activity) started with a time-frequency EEG analysis based on pure 3-cycles Morlet wavelets during a [-500 500] ms time interval and within a [6 50] Hz frequency window. The EEG baseline for the calculation of power was defined as the activity preceding the onset of the alerting central cue within the [−500 −300] ms time window (see Figure 1 for details on the timing of events). We performed two types of analyses: the first one concerned the frequency (30 Hz) and time of interest (beginning of burst delivery to target onset) across electrodes; the second one focused on the electrode of interest (FC2, the closest to the stimulated right FEF region) across time and frequencies. Direct planned comparisons (two-tailed paired t-tests at p<0.01) between active and sham trials separately for both rhythmic and random TMS patterns, and direct comparisons between rhythmic and random trials separately for both active and sham TMS were performed. For the first analysis, topographical maps of power and inter-trial coherence (i.e., topography of the scalp distribution of the power and inter-trial coherence) at specific frequency and time window of interest (30 Hz and [-133 0] ms; 0 being the target onset) were compared with paired t-tests calculated for every electrode. Similarly, for the second analysis, power across time and frequencies aligned to the target onset (commonly referred to as event-related spectrum perturbation, ERSP) and inter-trial coherence (ITC) at the electrode FC2 were compared with paired t-tests calculated for every time-frequency point during a restricted time window of interest [-300 200] ms. To correct for multiple direct planned comparisons, electrodes or time-frequency points that reached significance in the paired t-test were clustered and a non-parametric permutation test was applied on these clusters (10000 permutations, alpha = 0.05, (Maris and Oostenveld, 2007; Oostenveld et al., 2011), see Figure 2 and Figure 3 for results, only significant results are displayed). Because EEG data is highly correlated in space and exhibits physiological effects that last over several time points (i.e. an effect is likely to be spread over adjacent sensors and consecutive time points) cluster-based permutations is a highly sensitive method for solving the multiple comparison problem in this data (Maris and Oostenveld, 2007). However, in the case of a factorial design, there is no consensus on how to permute the data to correctly control for multiple comparisons when evaluating interaction effects between multiple factors (Suckling and Bullmore, 2004; Edgington and Onghena, 2007). For this reason, and driven by hypotheses of a different effect of rhythmic and random stimulation patterns on oscillatory activity, we chose to compute direct pairwise comparisons between our conditions.

**Figure 2.**
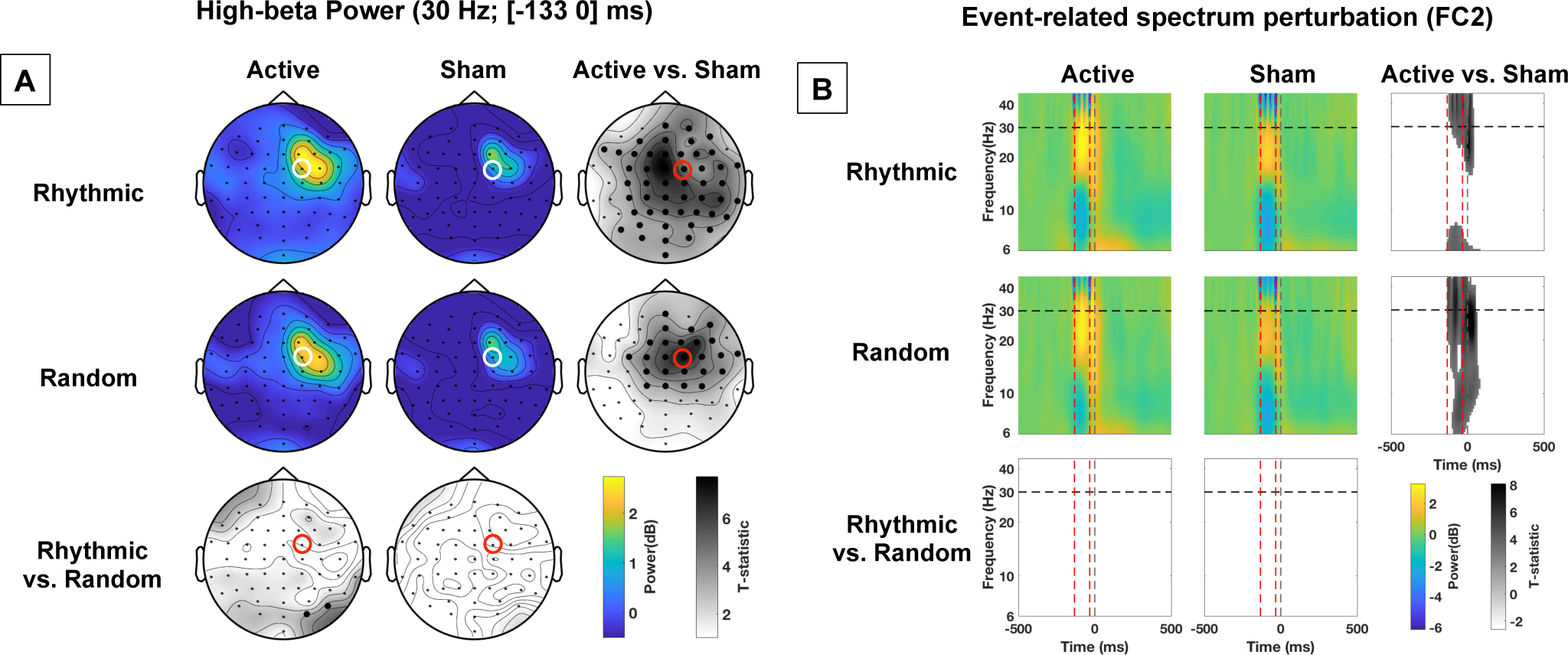
Impact of active vs. sham, rhythmic vs. random, right FEF stimulation on power. **(A)** Topographical maps representing power at 30 Hz during the [-133 0] ms pre-target onset time-window. The location of electrode FC2 (closest to the stimulated right FEF site), is indicated with a red or white open circle. On the statistical maps, electrodes from the topographic views for which EEG signals proved significantly different between conditions are signaled with bold black dots. Notice the increase of 30 Hz power for active vs. sham stimulation. **(B**) Event-related spectrum perturbation (ERSP) calculated for scalp electrode FC2. Red dotted vertical lines signal the onset and offset of the 4-pulse stimulation burst; the grey dotted vertical line signals the visual target onset. The horizontal black dotted line signals the frequency of 30 Hz at which active rhythmic patterns were delivered. On the statistical maps, grey colors signal statistically significant difference between conditions. Notice that both active rhythmic and active random stimulation increased power at ∼30 Hz during stimulation.

**Figure 3.**
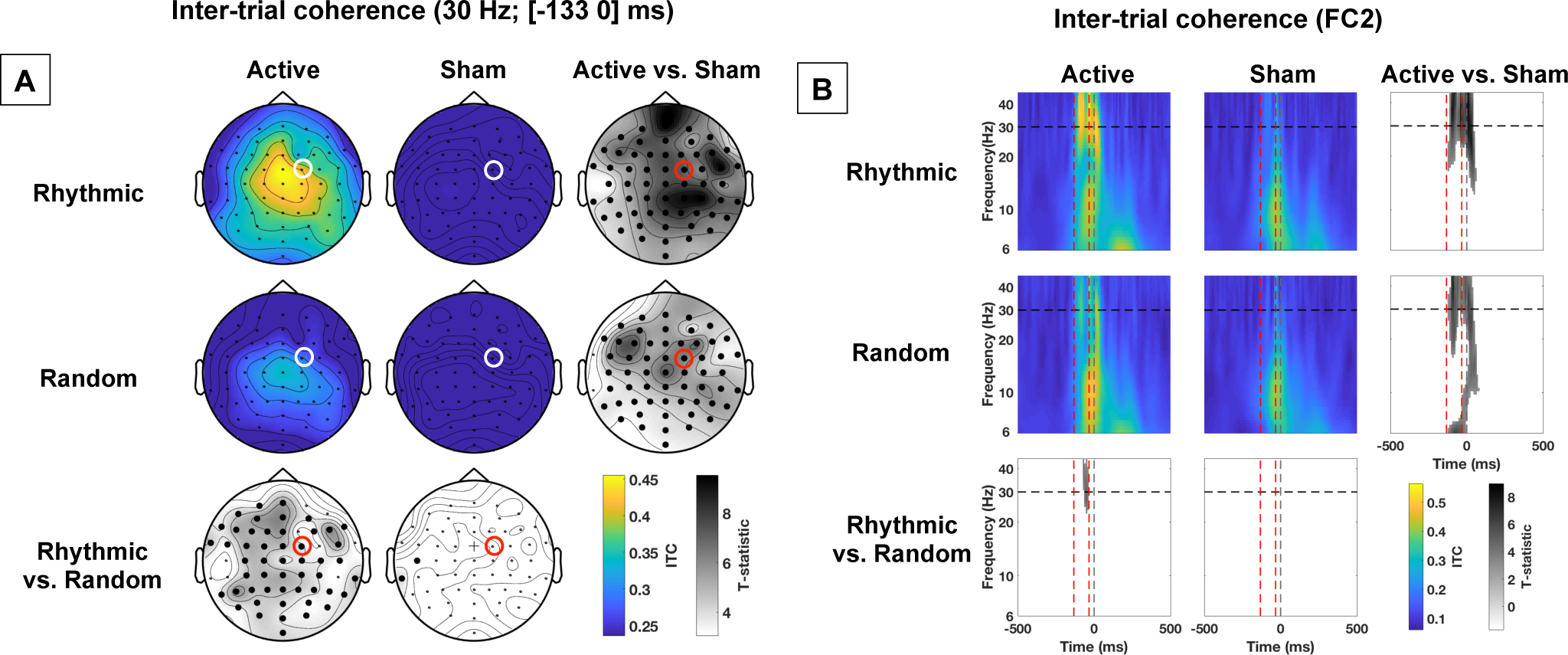
Impact of active vs. sham, rhythmic vs. random, right FEF stimulation on phase alignment (inter-trial coherence). **(A)** Topographical maps representing inter-trial coherence (ITC) of 30 Hz EEG activity during the [-133 0] ms pre-target onset time window. The location of electrode FC2 (closest to the stimulated right FEF site), is indicated with a red or white open circle. On the statistical maps, electrodes from the topographic views for which EEG signals proved significantly different between conditions are signaled with bold black dots; cross signs indicate marginally significant differences. **(B)** Inter-trial coherence (ITC) at scalp electrode FC2 throughout frequency bands and time windows. Red dotted vertical lines signal the onset and offset of the 4-pulse stimulation burst; the grey dotted vertical line signals the visual target onset. The horizontal black dotted line signals the frequency of 30 Hz at which active rhythmic patterns were delivered. On the statistical maps, grey colors signal statistically significant difference between conditions. Notice that a direct comparison between active *rhythmic* and *random* stimulation shows higher ITC for active *rhythmic* stimulation.

The ensuing procedure estimated evoked oscillations, i.e., high-beta EEG signals time-locked to the target onset and consequently to the TMS pulses (all of the pulses for rhythmic pattern, and the first and last pulses for the random pattern (Tallon-Baudry and Bertrand, 1999)). It followed the same rational as the calculation of ITC used to estimate phase alignment mentioned above. However, it allowed a finer analysis of our EEG data across different time windows before, during, and after the delivery of TMS bursts. Data were filtered ([25 35] Hz, see Figure 4 for results). Then, four time-windows of interest were defined as follows: T1: Pre TMS [−199.5 −133] ms; T2: TMS burst 1^st^ half (post pulses 1&2 for rhythmic TMS patterns) [−133 −66.5] ms; T3: burst 2^nd^ half (post pulses 3&4 for rhythmic TMS patterns) [-66.5 0] ms; T4: Visual Target [0 66.5] ms. The amplitude of evoked oscillations, was calculated by averaging the filtered data across trials, before averaging within each time window. The resulting amplitudes were analyzed using a trends-based repeated measures ANOVA with *stimulation pattern* (rhythmic, random), *stimulation condition* (active, sham) and *time window* (both linear and quadratic coefficients) as within-participant factors (see Figure 4C; note similar standard errors across conditions).

**Figure 4.**
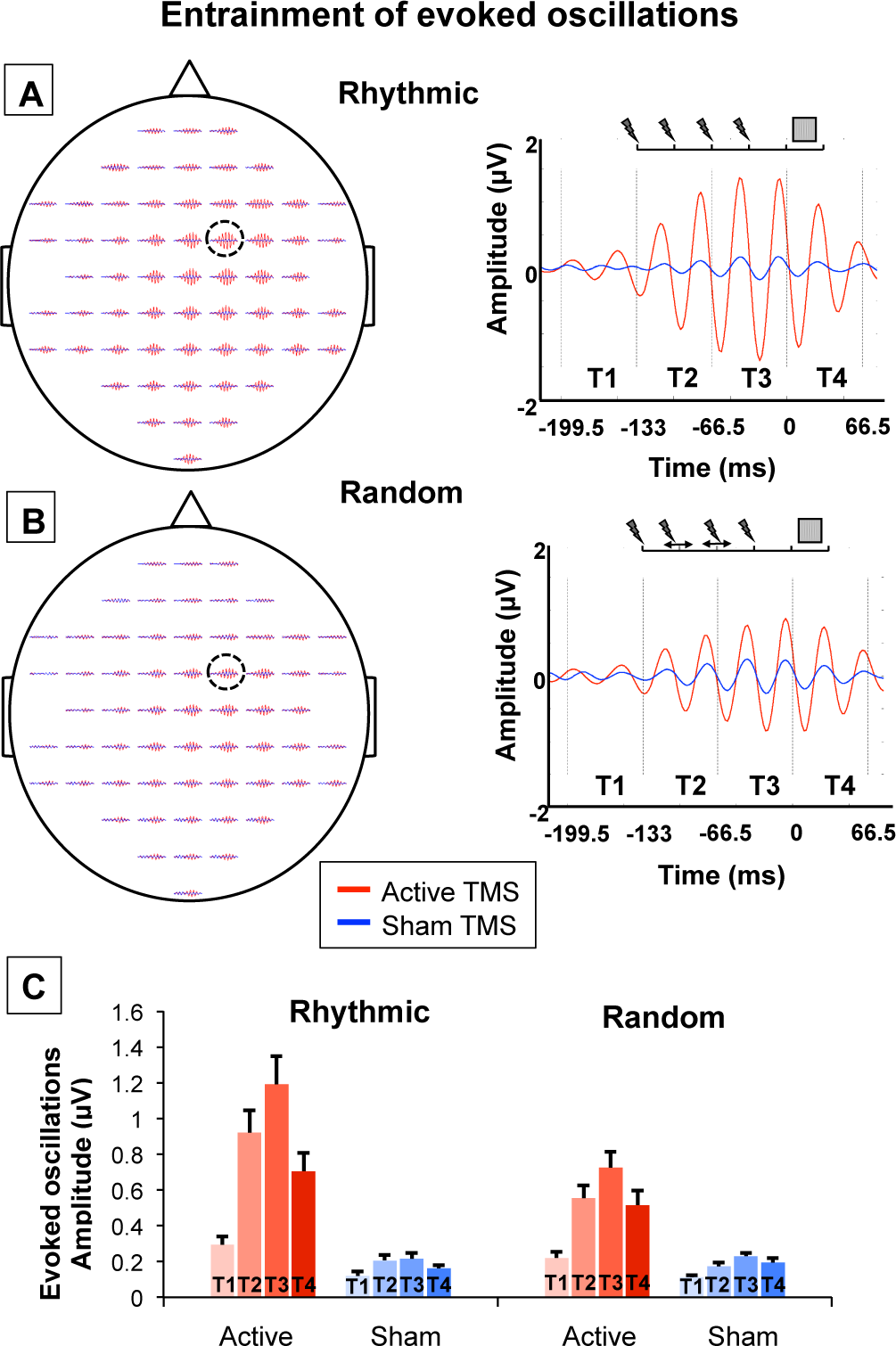
Causal impact of right FEF stimulation on evoked high-beta oscillations. Evoked oscillations (25-35 Hz, [-199.5 66.5] ms, [-2 2] µV) for rhythmic **(A)** and random **(B)** active/sham stimulation patterns for each of the 60 EEG scalp electrodes (left column; the location of electrode FC2, i.e., the closest to the stimulated right FEF, is indicated with an open circle) and at FC2 (right column). Vertical black dotted lines delineate the epochs employed for the analyses (T1: Pre TMS, T2: TMS burst part 1; T3: TMS burst part 2 and T4: Visual Target). Blue and red colors respectively represent the *sham* and *active* TMS conditions. Notice progressive increases in the amplitude of high-beta evoked oscillations (25-35 Hz), reaching higher levels during *rhythmic* than *random* active patterns throughout the course of 4-pulse stimulation patterns followed by a rather abrupt decay following the end of the burst. **(C)** Amplitude (mean and standard error) of evoked oscillations (25-35 Hz) for *rhythmic* and *random* active/sham stimulation patterns across the 4 time-windows of interest (T1: Pre TMS; T2: TMS burst part 1; T3: TMS burst part 2; and T4: Visual Target). Due to the complexity of representation of interaction effects, significant statistical results are not shown on the figure. Notice, however, that active *rhythmic* patterns caused higher amplitude increases of evoked oscillations than active *random* patterns (significant *stimulation pattern x stimulation condition* interaction). In addition, we found a progressive build-up of evoked oscillations along the course of the 4-pulse burst (amplitude T1<T2<T3), and a decay following the offset of the stimulation (T4<T3, significant *stimulation condition* x *time window* interaction).

In line with earlier findings and the most current mechanistic understanding of stimulation-induced entrainment (Thut et al., 2011a; Thut et al., 2017), the entrainment phenomenon occurs mainly in a frequency band centered around the stimulation frequency (Thut et al., 2011b; Amengual et al., 2017). On that basis, and also building on our prior experience in the domain (Chanes et al., 2013; Chanes et al., 2015; Quentin et al., 2015; Quentin et al., 2016), our task design and our analytical and statistical strategy were directed to assess changes in oscillatory activity centered in the TMS frequency (30 Hz) delivered to the right FEF. Specifically, we did not quantify higher gamma activity, potentially contaminated by muscles artifacts and line noise. Similarly, modulation of frequencies lower than mid-alpha, which developed at much longer time scale than the duration of the TMS burst and the time temporal window of interest chosen for our analyses, was not statistically assessed. For these reasons, we cannot rule out the possibility that 30 Hz FEF TMS also modulated oscillatory activity outside of the high-beta range.

### Behavioral data analyses

Visual detection performance was assessed with perceptual sensitivity (d’) and response criterion (β). These two outcome measures are employed in Signal Detection Theory (SDT) to characterize detection performance when it can be strongly influenced by belief (e.g., when stimuli are presented around the perceptual threshold) (Stanislaw and Todorov, 1999). Perceptual sensitivity is a bias-free measure of the participants’ ability to detect a target, whereas response criterion describes the relative preference (bias) of participants for one response over the alternative one in case of doubt (i.e., in our case, a preference for ‘yes, I saw the target’ over ‘no, I did not see it’).

To compute these measures, trials in which the location of the target was correctly reported by participants were considered correct detections or “hits”; trials in which the presence of the target was not acknowledged were considered “misses”; trials in which participants reported a location for a target that was not present were considered “false alarms”; trials in which the target was absent and participants correctly reported to not have seen it were considered “correct rejections”. Trials in which the location of a present target was incorrectly reported were counted as “errors” and were excluded from the main analyses (given the impossibility to distinguish whether participants incorrectly detected the target or correctly detected it but pressed the wrong button). Following an established procedure, zero false alarms were replaced by half false alarms (0.5) in order to calculate d’ and β measures (Macmillan and Creelman, 2005). Perceptual sensitivity and response bias were calculated as follows: d’=ϕ^−1^(H)-ϕ^−1^(FA) and β=N(ϕ^−1^(H))/N(ϕ^−1^(FA)), where ϕ^−1^ is the z-transform, N the un-cumulated density function, H the hit rate, and FA the false alarm rate.

For statistical analyses, d’ and β were each subjected to a 2×2×2 repeated-measure ANOVA with *stimulation pattern* (rhythmic, random), *stimulation condition* (active, sham) and *target location* (left, right) as within-participant factors. Pairwise comparisons between specific conditions were performed with t-student tests. The ANOVA on response criterion (β) did not reveal any significant main effect or any significant interaction (all the p>0.11). Thus, only effects on perceptual sensitivity (d’) are presented in the main text of the manuscript (see also Figure 5A; note similar standard errors across conditions).

**Figure 5.**
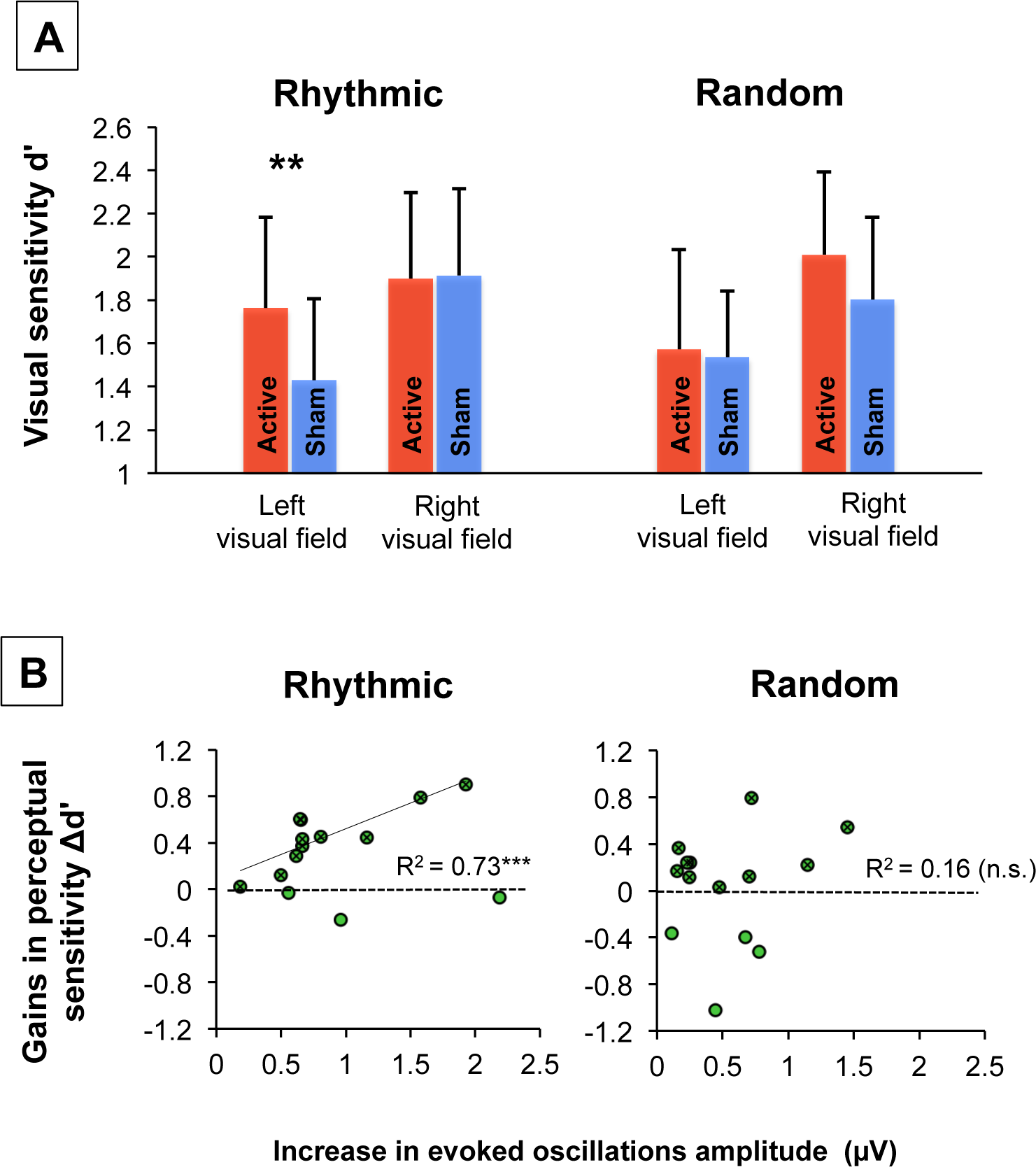
Causal impact of right FEF stimulation on conscious visual detection and relationship with entrained high-beta oscillations. **(A)** Group impact of active/sham rhythmic and random patterns delivered to the right FEF on the conscious detection performance of near-threshold targets presented in the left or right visual fields (means and standard errors; statistical comparison: **p<0.01). Importantly *rhythmic* (but not *random)* right FEF active stimulation which, according to EEG evidence (see Figures 1 and 2), increased high-beta power and inter-trial coherence, also increased visual sensitivity (d’) for targets displayed in the left visual hemifield. **(B)** Correlation plots between levels of high-beta entrainment (estimated through increases of amplitude of evoked oscillations between active and sham TMS) and conscious visual performance gains (d’ active TMS - d’ sham TMS) with *rhythmic* (left) or *random* (right) active TMS patterns for targets presented in the left visual field. Green dots represent all participants (n=14). Crossed green dots represent pools of participants (n=11 for high-beta rhythmic TMS, n=10 for random TMS) who experienced visual sensitivity (d’) increases with right FEF stimulation. For high-beta rhythmic TMS, a linear correlation with only the latter selected cohort of participants (green regression line) proved highly significant, whereas for random TMS, no correlation reached significance (***p<0.001; n.s. non-significant).

Correlations between significant rhythmic or random TMS-driven (active-sham) modulation of evoked oscillations, and rhythmic or random TMS-driven (active-sham) modulation of conscious visual sensitivity (d’), were tested with a Pearson’s correlation coefficient test.

### Further statistical considerations

Statistical significance was set at *p*<0.05 and tests were two-tailed. To further strengthen the validity of our findings, we indicated the 95%-Confidence Intervals (CI) and Cohen’s d effect size (interpretation: d=0.2: small; d=0.5: medium; d=0.8: large effect size (Cohen, 1988)) when appropriate. To further test our main conclusions, we also calculated Bayes factors. For this, we used a uniform interval bound by the compared values (with opposite signs); when necessary, we corrected for the number of degrees of freedom as indicated by Dienes (2014) (interpretation: B<0.3: substantial evidence for the null hypothesis; B>3: substantial evidence for the alternative hypothesis). Finally, we performed bootstrap statistics based on 1000 correlation coefficients calculated on datasets drawn from the original dataset with replacement (alpha=0.05). To compare the strength of different correlations, we calculated unpaired t-tests between the Bootstrap distributions of correlation coefficients and the Z_2_* as indicated in Steiger (1980).

## Results

Our group of healthy participants performed a visual detection task, where they had to report if they detected a laterally presented target or not and, if they did, where in the screen it appeared. Within the same blocks of trials, active or sham TMS was delivered to the right FEF shortly before the onset of the visual target (see Fig. 1A and 1B). Different patterns of TMS were delivered in separate blocks: rhythmic (30 Hz) or random (no specific frequency) bursts made of 4 consecutive TMS pulse for a total duration of 100 ms (see Fig. 1C). EEG was continuously recorded during the task.

### Impact of high-beta right frontal stimulation on event-related spectrum perturbation

Event-related spectrum perturbation (ERSP) analyses of our EEG data assessed modulations of power across different time-frequency points and electrodes. We first examined the modulation of power for all electrodes at a specific frequency (30 Hz) and time window ([-133 0] ms; 0 being the target onset; see Figure 2A) of interest. Second, we examined the modulation of power at the electrode of interest, i.e., the closest to the stimulated right FEF (FC2), for a broader frequency range ([6 45] Hz) and time window ([-500 500] ms, see Figure 2B).

The first analysis revealed that both *rhythmic* and *random* active right FEF stimulation significantly increased high-beta power at 30 Hz, compared to their sham controls. This increase was rather widespread, reaching significance in scalp electrodes over frontal, central, and parietal sites for the active vs. sham *rhythmic* TMS conditions. In contrast, the active vs. sham *random* TMS conditions comparison showed a more spatially restricted impact centered over right frontal sites (see Figure 2A, the bold black dots on the Statistics maps indicate electrodes showing significant differences between active and sham TMS conditions). No consistent differences in 30 Hz power (with the exception of two isolated parietal electrodes) were found when directly comparing active *rhythmic* vs. active *random* stimulation. The second analysis revealed that high-beta power was significantly enhanced on the FC2 recordings during *rhythmic* stimulation (active vs. sham comparison) around 30 Hz (∼ [25-45] Hz frequency band), and during *random* stimulation (active vs. sham comparison) across the whole spectrum of tested frequencies ([6-45 Hz]) (see Figure 2B).

These analyses suggest that both rhythmic and random active TMS patterns (which only differed with regards to the temporal distribution of their 4 pulses) significantly enhanced high-beta power. The spatial and frequency signatures of these effects appear potentially distinct; however, such apparent differences did not reach statistical significance when *rhythmic* and *random* TMS were directly compared.

### Impact of rhythmic activity on the phase alignment of right frontal activity

The impact of stimulation on phase alignment was studied through the analysis of inter-trial coherence (ITC), a measure that assesses the consistency across trials of a neurophysiological signal phase. Data from either the complete electrode grid (30 Hz and [-133 0] ms) or specifically from the FC2 electrode (extended time-frequency window) revealed that both *rhythmic* and *random* stimulation significantly increased phase alignment at the high-beta band. Nonetheless, the direct comparisons between the two active conditions (which only differed in the temporal distribution of their 4 TMS pulses) showed higher ITC for the active *rhythmic* than for the active *random* stimulation condition (see Figure 3).

### Impact of rhythmic TMS activity on frontal evoked oscillations

Analyses assessing the magnitude of time-locked high-beta [25 35] Hz filtered EEG signals (i.e., evoked oscillations) aimed to further prove that active rhythmic 30 Hz patterns entrained local oscillations. To that end, we averaged the amplitude of evoked oscillations within 4 time-windows of interest (T1 to T4, see Figure 4 for details).

Figure 4 reveals progressive increases of the beta oscillations across the first 3 time windows (T1<T2<T3), followed by decay in the last time window (T3>T4). Such a quadratic trend appeared larger for active than sham conditions, and more so for rhythmic than random patterns. To evaluate the significance of these observations, a trends-based ANOVA with the factors *stimulation pattern, stimulation condition* and *time window*, the latter being evaluated by linear and quadratic coefficients, was used. The ANOVA showed no evidence for lack of fit of the model (p=0.69) and showed significant main effects of *stimulation pattern* (F(1,13)=15.26, p<0.001), *stimulation condition* (F(1,13)=177.42, p<0.0001) and *time window* (linear: F(1,13)=23.26, p<0.0001; quadratic: F(1,13)=46.10, p<0.0001). More importantly, this ANOVA also showed a significant *stimulation pattern* (rhythmic, random) x *stimulation condition* (active, sham) interaction (F(1,13)=15.69, p<0.001), revealing that active vs. sham increases in amplitude of evoked oscillatory activity were significantly higher for *rhythmic* than for *random* stimulation (p<0.0001, CI=[0.19 0.36], d=0.36, B_U[-0.30 0.36]_=105; both active vs. sham differences were highly significant when tested separately, p<0.0001; for rhythmic: d=0.94; for random: d=0.81). Furthermore, *stimulation condition* (active, sham) interacted with both the linear (F(1,13)= 11.57, p<0.001) and the quadratic coefficients (F(1,13)= 25.77, p<0.0001) of the time-window; the (positive) linear coefficient was higher for active than sham stimulation (CI=[0.60 1.31], d=0.84, B_U[-0.181.13]_=44), and the (negative) quadratic coefficient was lower for active than sham stimulation (CI=[−0.24 −0.10], d=0.77, B_U[-0.03 0.20]_=9745). The *stimulation pattern* (rhythmic, random) interacted with the quadratic coefficient (F(1,13)= 4.90, p<0.05); the (negative) quadratic coefficient was lower for rhythmic than random stimulation (CI=[−0.15 −0.01], d=0.28, B_U[-0.08 0.16]_=3.02). Finally, there was a trend towards significance for the 3-way interaction including the quadratic component (F(1,13)= 3.54, p=0.061), which could be explained by a larger active vs. sham decrease of the (negative) quadratic coefficient for rhythmic than for sham stimulation (p<0.05; CI=[−0.26 −0.03]; d=0.46; B_U[-0.10 0.24]_=2.26; both active vs. sham differences being significant when tested separately, p<0.01; for rhythmic: d=0.93; for random: d=0.73).

To summarize, evoked high-beta oscillatory activity entrained by rhythmic 30-Hz patterns prior to the onset of the visual target increased progressively across the 4-pulse TMS burst, and decayed rather rapidly after the offset of the stimulation pattern. Together with the increase of power and ITC, these results support the ability of *rhythmic* 30 Hz patterns to entrain higher levels of high-beta oscillations in right FEF regions than *random* patterns. This suggests that rhythmic stimulation is the optimal pattern to enhance power and increase phase alignment, i.e., to entrain *oscillatory* activity at the stimulation input frequency.

### Impact of right frontal rhythmic stimulation on conscious visual performance

The contiguous behavioral consequences of high-beta oscillation entrainment on conscious visual detection performance (i.e., visual sensitivity d’) were explored with a 2×2×2 repeated-measure ANOVA with the factors: *stimulation pattern* (rhythmic, random), *stimulation condition* (active, sham) and *target location* (left, right). This ANOVA showed main effects of *stimulation condition* (real, sham: F(1,13)=5.33; p<0.05) and *target location* (right, left: F(1,13)=10.14; p<0.01), supporting higher conscious visual detection performance under active than sham stimulation and also for targets displayed in the right than the left visual hemifield (Figure 5A).

A trend towards statistical significance for the triple interaction (F(1,13)=3.97; p=0.068) was found. Despite the fact that this interaction was only close to significance, a direct comparison is of high interest for testing our *a-priori* hypothesis that rhythmic and random effects on perception might be different (Chanes et al., 2013; Chanes et al., 2015; Quentin et al., 2016). Thus, we performed t-tests, which showed that rhythmic activity increased visual sensitivity (d’) for targets presented in the left, i.e. contralateral to the stimulation (p<0.005, CI=[0.14 0.53], d=0.42), but not in the right visual hemifield (p>0.88, CI=[-0.23 0.20], d=0.02). In contrast, random stimulation failed to influence visual detection sensitivity for targets in either of the two hemifields (both p>0.11 and d={0.05; 0.27}) (see Figure 5A). For the left visual field, the Bayes factor fell short of the conventional criteria of substantial evidence of higher increase following rhythmic than random pattern (B_U[-0.037 0.332]_=2.63) whereas for right visual field, the evidence was less conclusive (B_U[-0.014 0.208]_=1.81).

Our analyses suggest that brief *rhythmic* stimulation patterns entrained high-beta activity in the right FEF prior to the onset of a lateralized visual target, and that such entrained oscillatory activity is causally related to the improvement conscious detection performance for targets displayed in the left visual hemifield.

### Correlations between entrainment and conscious visual performance gains

Aiming to provide additional support for a causal link between the entrainment of right frontal high-beta oscillations and improvements of conscious visual detection, we correlated the outcome measure gauging levels of entrainment (i.e., active–sham rhythmic TMS differences in the amplitude of evoked oscillations for time window T3) and gains of conscious visual detection performance (i.e., active–sham rhythmic TMS visual sensitivity (d’) differences for left hemifield targets).

When all participants were included in the analysis, the Pearson’s correlation coefficient between these two measures failed to reach significance (R^2^=0.06; p=0.39; df=12; p_bootstrap_=0.90). Nonetheless, given this null result, we formulated another hypothesis relying on the basis that inter-individual differences in the structural connectivity involving the FEF has often explained the failure of right frontal stimulation to modulate visual performance (Quentin et al., 2013; Quentin et al., 2015; Quentin et al., 2016). Thus, we computed the same analysis with the 11 participants showing increases of visual sensitivity with 30 Hz rhythmic stimulation. It revealed a highly significant correlation between these two variables (R^2^=0.72; p<0.001; df=9, Figure 5B left; p<0.05 after Bonferroni correction; p_bootstrap_<0.05). These results show that when rhythmic stimulation enhanced conscious visual perception, the magnitude of the evoked oscillation entrainment correlated significantly with increases in visual performance for left targets.

Several observations strengthen the specificity of this significant correlation between evoked oscillations entrained by rhythmic TMS patterns and behavioral outcomes. First, we failed to find a significant correlation between the same variables for *random* stimulation patterns, neither when considering all participants, nor when selecting the 11 participants mentioned above or a newly selected cohort of participants attesting increases of visual sensitivity for left targets under *random* stimulation (all 3 analyses p>0.25 and p_bootstrap_>0.25). Second, the significant correlation coefficient for active *rhythmic* stimulation with the selection of 11 participants proved significantly higher than the non-significant correlation coefficient shown for active *random* stimulation (p<0.0001; Z_2_*=2.27). Third, none of the correlations between increases of evoked oscillations (active-sham) for time-window T3 and increase of visual sensitivity (active-sham) for right targets (instead of left targets) proved significant (p>0.05 and p_bootstrap_>0.20 for both *rhythmic* and *random* stimulation conditions, regardless of the selection of participants considered). Moreover, the significant correlation coefficient for *rhythmic* pattern and visual sensitivity gains for left targets was significantly higher than the non-significant correlation coefficient for right targets (p<0.0001; Z_2_*=2.86 for rhythmic patterns and p<0.0001; Z_2_*=2.47 for *random* patterns).

## Discussion

Our data support the notion that the entrainment of high-beta oscillations in the right FEF, shortly before the onset of a low-contrast target, causally increases conscious visual detection performance. Electrophysiological (EEG) recordings revealed that rhythmic stimulation entrained oscillations at the input frequency band (∼30 Hz). Indeed, both *rhythmic* and *random* patterns increased high-beta power. Nonetheless, *rhythmic* patterns induced significantly higher levels of phase alignment than *random* activity. Moreover, the amplitude of evoked oscillations, phase-aligned to the stimulation, built up during the course of a 4-pulse stimulation burst, and achieved significantly higher amplitude for *rhythmic* than for *random* stimulation. It then decayed rather rapidly after its offset. These outcomes support the ability of rhythmic TMS to noninvasively manipulate local synchrony in circumscribed cortical regions and impose specific patterns of oscillatory activity, which might serve to explore, enhance or even restore human behaviors in the near future.

Among the main techniques currently available to manipulate oscillatory activity in humans, namely rhythmic TMS (Klimesch et al., 2003; Romei et al., 2010; Thut and Pascual-Leone, 2010; Thut et al., 2011a; Thut et al., 2011b; Romei et al., 2012; Chanes et al., 2013; Hanslmayr et al., 2014; Ruzzoli and Soto-Faraco, 2014; Quentin et al., 2015; Quentin et al., 2016) and tACS (Kanai et al., 2008; Zaehle et al., 2010; Antal and Paulus, 2013; Brittain et al., 2013; Helfrich et al., 2014b; Helfrich et al., 2014a), we opted for the former, given its higher focality and special ability to entrain on a trial-by-trial basis episodic events of oscillatory activity during specific time-windows, which is crucial for probing focal contributions to ongoing human cognitive processes and behaviors. Transcranial ACS remains better suited to induce interregional synchronization across large cortical areas and explore research questions for which the timing of oscillatory events with regards to behavioral correlates is not critical.

Entrainment of biological rhythms emerges from a progressive phase alignment of different local oscillators following the rhythms dictated by either internal, or by external, “pacemakers”, which in our study were provided by focal rhythmic TMS (Thut et al., 2011a; Thut et al., 2011b; Thut et al., 2017). Consequently, simultaneous EEG recordings should show both increases in power at the input frequency and a progressive phase alignment.

The progressive build-up of high-beta oscillations during the course of the burst featured in our data grants convincing support in favor of rhythmic entrainment. It disentangles the synchronization of neural assemblies at the stimulation frequency from a mere injection of rhythmic activity arising from evoked potentials triggered by each individual TMS pulse (Komssi et al., 2004; Bonato et al., 2006). Indeed, whereas evoked potentials generated by individual pulses tend to keep a similar amplitude, increases of post-pulse activity throughout the course of the burst (a measure that in our study proved significantly higher during *rhythmic* than *random* active TMS patterns) are most likely associated with a phase alignment of local cortical oscillators.

Taking these criteria into account, our data indicate that while both active *rhythmic* and *random* stimulation patterns yielded power increases within the high-beta band compared to their associated sham bursts (see statistical maps Figure 2), phase alignment was higher for rhythmic than random stimulation (see statistical maps Figure 3). Consequently, 30 Hz *rhythmic* patterns led to superior increases in the amplitude of evoked high-beta oscillations and resulted in stronger entrainment of beta rhythmic activity (see Figure 4). Analogously, *rhythmic* but not *random* active TMS patterns increased visual sensitivity for the detection of targets presented in the contralateral hemifield (see Figure 5A), a result that replicates prior observations made by our own group under similar conditions (Chanes et al., 2013; Quentin et al., 2015; Quentin et al., 2016). Additionally, in participants who showed enhanced visual sensitivity with rhythmic stimulation, the entrainment of high-beta oscillations scaled with visual performance increases (see Figure 5B). This strengthens the evidence in favor of a causal role for high-beta right frontal rhythms in driving access to visual consciousness.

Pioneering evidence in the field of rhythmic non-invasive stimulation has suggested that the entrainment of oscillations results from the alignment of local oscillators operating at their so-called “natural” frequency (Thut et al., 2011b) or in neighboring oscillation bands (Klimesch et al., 2003). Since brain areas tend to naturally oscillate within specific frequency ranges (Rosanova et al., 2009), local synchrony within a given cortical location cannot be easily imposed at any frequency. In our study, the successful entrainment of high-beta oscillations in the human right FEF, a region of the fronto-parietal network that synchronizes at this same frequency range during the allocation of endogenous spatial attention (Buschman and Miller, 2007; Phillips and Takeda, 2009), supports this view. Indeed, developing extensive knowledge on the status of local and network physiological brain activity, and most particularly features of local and interregional synchrony recorded via EEG, is part of the recently established *information-based approach* to non-invasive stimulation, aiming to guide the selection of TMS parameters and optimize the use of these tools in exploratory or applied clinical settings (Romei et al., 2016a).

Prior evidences in favor of high-beta power increases over both right frontal and parietal areas also suggest that oscillation entrainment in the FEF spreads to interconnected areas across a right-lateralized fronto-parietal network (Quentin et al., 2015; Quentin et al., 2016). Therefore, the improvement of visual performance reported as causally associated to high-beta oscillations is likely to be mediated by an engagement of top-down attention subtended by the dorsal attentional network (Corbetta et al., 2008; Vernet et al., 2014b) that synchronizes within this same band increase during top-down attention processes (Buschman and Miller, 2007; Phillips and Takeda, 2009). Such improvement would be related to the demonstrated ability of attentional orienting mechanisms to facilitate the detection of lateralized near-threshold visual targets (Carrasco, 2011).

Although top-down attention is a selective mechanism often leading to higher visual awareness (Posner, 1994), the two processes can be dissociated (Lamme, 2003). Examining the interactions between attention and awareness requires the direct manipulation of attention, e.g., by using spatial cues to orient attention to an area of visual space prior to target onset (Grosbras and Paus, 2002; Chanes et al., 2012). For methodological reasons (essentially the large number of trials needed per conditions for meaningful TMS-EEG analyses), our behavioral paradigm did not directly manipulate visuospatial attention. Moreover, participants were not requested to answer as fast as possible (preventing the analysis of reaction times, which are often used as a proxy for attentional orienting). Hence, on the basis of our data, we can neither confirm nor exclude that the reported effects on right frontal high-beta oscillations and visual awareness were subtended by an attentional mechanism. Investigating the specific role of attention in the causal influence of FEF beta activity on visual consciousness will require a specific design and will have to be addressed in future ad hoc experiments.

We would like to emphasize that the possibility to entrain beta oscillations directly by stimulating the right FEF does not necessarily imply that natural beta oscillations are exclusively of cortical origin. Indeed, thalamo-cortical loops have been shown to play a role in both the regulation of cortical oscillations and attentional and awareness processes (Saalmann and Kastner, 2011). Furthermore, such loops might be regulated by dopamine release in the basal ganglia, which has been in turn associated to beta oscillations (13-30 Hz) in motor networks (Jenkinson and Brown, 2011) and to improvements in subjective and objective visual performance (Van Opstal et al., 2014). The extent to which the level of dopamine and/or ongoing beta oscillations is influencing the magnitude of TMS-evoked oscillations and the increase of perceptual sensitivity was out of the scope of the current study, but remains an interesting topic of investigation for future studies.

Single-pulse TMS over the right FEF has also been shown to speed-up discrimination and/or increase detection and visual awareness (Grosbras and Paus, 2002, 2003; Chanes et al., 2012). This evidence suggests that the enhancement of visual performance and awareness could also result from TMS-driven increases of background activity, drifting closer to threshold, hence helping weak signals to reach perceptual threshold and become consciously visible (Grosbras and Paus, 2003). Alternatively, enhancement could have resulted from boosting only specific clusters of neurons according to their level of activity (O’Shea and Walsh, 2004). Such effects could result from local activation within the FEF, or from top-down modulations of occipital brain regions, which enhance the gain of incoming visual signals (Grosbras and Paus, 2003). To this regard, single TMS pulses delivered to the right FEF have shown to modulate occipital excitability measured with phosphene threshold (Silvanto et al., 2006). Similarly, short rTMS bursts (at ∼10 Hz) to the FEF modulated both visual evoked potentials (Taylor et al., 2007) and fMRI BOLD responses (Ruff et al., 2006) in occipital areas. Finally, single TMS pulses delivered to a frontal area close to the FEF have shown to enhance the power of the so-called “natural” beta band activity characteristic from the stimulated region (Rosanova et al., 2009).

Whether or not the perceptual effects of single-pulse TMS are directly or indirectly related to the increase of beta (∼30 Hz) oscillations remains to be elucidated. Conversely, it is possible that the conscious visual sensitivity increase reported in our study might not be solely contributed by oscillatory entrainment mechanisms. However, the significantly higher behavioral performance and high-beta entrainment driven by rhythmic, as compared to random, TMS patterns, and the correlation between the amplitude of evoked oscillatory and perceptual facilitation, strongly support the causal contribution of right frontal oscillations to visual awareness.

In conclusion, our study provides support to our ability to entrain episodes of local synchrony subtending a specific cognitive function. This finding provides causal evidence in humans that oscillatory activity (in this study, high-beta at 30 Hz), operating focally within a cortical region (the right FEF), contributes to a cognitive processing and behavior (access to consciousness and visual detection performance). Furthermore, our results support the use of well-established invasive (e.g. intracranial or deep brain stimulation) and noninvasive (rhythmic TMS or tACS) brain stimulation technologies and also emerging experimental approaches (e.g., optogenetic or ultrasound stimulation) to manipulate cortical synchronization. This opens new avenues to explore, improve and restore behaviors subtended by focal dysfunctions of brain synchrony.

## Acknowledgement

Marine Vernet and Romain Quentin were supported by fellowships from the *Fondation pour la Recherche Medicale*. Chloé Stengel was supported by a PhD fellowship from the University Pierre and Marie Curie. Julià L. Amengual was supported by a fellowship from the *Fondation Fyssen*. The activities of the laboratory of Dr. Valero-Cabré are supported by research grants IHU-A-ICM-Translationnel, ANR projet Générique OSCILOSCOPUS and Flag-era-JTC-2017 CAUSALTOMICS. The authors would also like to thank Juliette Godard and Laura Fernandez for providing help during data acquisition, Sara Ahmed for final proof reading of the manuscript, and the Naturalia & Biologia Foundation for financial assistance for traveling and attending meetings.

## Author contributions

Conceptualization: M.V., R.Q. and A.V.C. Data acquisition: M.V., R.Q. Data analysis: M.V. and C.S. Manuscript preparation: M.V., C.S., R.Q., J.L.A. & A.V.C. Supervision: A.V.C.

## Data availability statement

Data are available from the corresponding author upon request.

## References

Amengual JL, Vernet M, Adam C, Valero-Cabre A (2017) Local entrainment of oscillatory activity induced by direct brain stimulation in humans. Sci Rep 7:41908.

Antal A, Paulus W (2013) Transcranial alternating current stimulation (tACS). Front Hum Neurosci 7:317.

Bonato C, Miniussi C, Rossini PM (2006) Transcranial magnetic stimulation and cortical evoked potentials: a TMS/EEG co-registration study. Clin Neurophysiol 117:1699–1707.

Brainard DH (1997) The Psychophysics Toolbox. Spat Vis 10:433–436.

Brittain JS, Probert-Smith P, Aziz TZ, Brown P (2013) Tremor suppression by rhythmic transcranial current stimulation. Curr Biol 23:436–440.

Buschman TJ, Miller EK (2007) Top-down versus bottom-up control of attention in the prefrontal and posterior parietal cortices. Science 315:1860–1862.

Buzsaki G, Draguhn A (2004) Neuronal oscillations in cortical networks. Science 304:1926–1929.

Carrasco M (2011) Visual attention: the past 25 years. Vision research 51:1484–1525.

Chanes L, Chica AB, Quentin R, Valero-Cabre A (2012) Manipulation of pre-target activity on the right frontal eye field enhances conscious visual perception in humans. PLoS One 7:e36232.

Chanes L, Quentin R, Tallon-Baudry C, Valero-Cabre A (2013) Causal Frequency-Specific Contributions of Frontal Spatiotemporal Patterns Induced by Non-Invasive Neurostimulation to Human Visual Performance. J Neurosci 33:5000–5005.

Chanes L, Quentin R, Vernet M, Valero-Cabre A (2015) Arrhythmic activity in the left frontal eye field facilitates conscious visual perception in humans. Cortex 71:240–247.

Cohen J (1988) Statistical power analysis for the behavioral sciences, 2nd Edition. Hillsdale, N.J.: L. Erlbaum Associates.

Corbetta M, Patel G, Shulman GL (2008) The reorienting system of the human brain: from environment to theory of mind. Neuron 58:306–324.

Cornsweet TN (1962) The Staircase-Method in Psychophysics. American J Psychol 75:485–491.

Delorme A, Makeig S (2004) EEGLAB: an open source toolbox for analysis of single-trial EEG dynamics including independent component analysis. Journal of neuroscience methods 134:9–21.

Dugue L, VanRullen R (2017) Transcranial Magnetic Stimulation Reveals Intrinsic Perceptual and Attentional Rhythms. Front Neurosci 11:154.

Edgington E, Onghena P (2007) Randomization tests: Chapman and Hall/CRC.

Fiebelkorn IC, Pinsk MA, Kastner S (2018) A Dynamic Interplay within the Frontoparietal Network Underlies Rhythmic Spatial Attention. Neuron 99:842–853 e848.

Fries P, Nikolic D, Singer W (2007) The gamma cycle. Trends Neurosci 30:309–316.

Fuggetta G, Pavone EF, Walsh V, Kiss M, Eimer M (2006) Cortico-cortical interactions in spatial attention: A combined ERP/TMS study. J Neurophysiol 95:3277–3280.

Grosbras MH, Paus T (2002) Transcranial magnetic stimulation of the human frontal eye field: effects on visual perception and attention. J Cogn Neurosci 14:1109–1120.

Grosbras MH, Paus T (2003) Transcranial magnetic stimulation of the human frontal eye field facilitates visual awareness. Eur J Neurosci 18:3121–3126.

Hamidi M, Slagter HA, Tononi G, Postle BR (2010) Brain responses evoked by high-frequency repetitive TMS: An ERP study. Brain Stimulat 3:2–17.

Hanslmayr S, Matuschek J, Fellner MC (2014) Entrainment of prefrontal beta oscillations induces an endogenous echo and impairs memory formation. Curr Biol 24:904–909.

Helfrich RF, Schneider TR, Rach S, Trautmann-Lengsfeld SA, Engel AK, Herrmann CS (2014a) Entrainment of brain oscillations by transcranial alternating current stimulation. Current biology: CB 24:333–339.

Helfrich RF, Knepper H, Nolte G, Struber D, Rach S, Herrmann CS, Schneider TR, Engel AK (2014b) Selective modulation of interhemispheric functional connectivity by HD-tACS shapes perception. PLoS Biol 12:e1002031.

Jenkinson N, Brown P (2011) New insights into the relationship between dopamine, beta oscillations and motor function. Trends Neurosci 34:611–618.

Kahkonen S, Komssi S, Wilenius J, Ilmoniemi RJ (2005) Prefrontal TMS produces smaller EEG responses than motor-cortex TMS: implications for rTMS treatment in depression. Psychopharmacology (Berl) 181:16–20.

Kanai R, Chaieb L, Antal A, Walsh V, Paulus W (2008) Frequency-dependent electrical stimulation of the visual cortex. Curr Biol 18:1839–1843.

Klimesch W, Sauseng P, Gerloff C (2003) Enhancing cognitive performance with repetitive transcranial magnetic stimulation at human individual alpha frequency. Eur J Neurosci 17:1129–1133.

Komssi S, Kahkonen S, Ilmoniemi RJ (2004) The effect of stimulus intensity on brain responses evoked by transcranial magnetic stimulation. Hum Brain Mapp 21:154–164.

Lamme VA (2003) Why visual attention and awareness are different. Trends Cogn Sci 7:12–18.

Macmillan N, Creelman C (2005) Detection theory: a user’s guide. London: Erlbaum Associates.

Maris E, Oostenveld R (2007) Nonparametric statistical testing of EEG- and MEG-data. J Neurosci Methods 164:177–190.

Marshall TR, Bergmann TO, Jensen O (2015) Frontoparietal Structural Connectivity Mediates the Top-Down Control of Neuronal Synchronization Associated with Selective Attention. PLoS Biol 13:e1002272.

Moore T, Fallah M (2001) Control of eye movements and spatial attention. Proc Natl Acad Sci U S A 98:1273–1276.

O’Shea J, Walsh V (2004) Visual awareness: the eye fields have it? Curr Biol 14:R279–281.

Oostenveld R, Fries P, Maris E, Schoffelen JM (2011) FieldTrip: Open source software for advanced analysis of MEG, EEG, and invasive electrophysiological data. Comput Intell Neurosci 2011:156869.

Pareti G, De Palma A (2004) Does the brain oscillate? The dispute on neuronal synchronization. Neurol Sci 25:41–47.

Paus T (1996) Location and function of the human frontal eye-field: a selective review. Neuropsychologia 34:475–483.

Phillips S, Takeda Y (2009) Greater frontal-parietal synchrony at low gamma-band frequencies for inefficient than efficient visual search in human EEG. Int J Psychophysiol 73:350–354.

Posner MI (1994) Attention: the mechanisms of consciousness. Proc Natl Acad Sci U S A 91:7398–7403.

Quentin R, Chanes L, Vernet M, Valero-Cabre A (2015) Fronto-Parietal Anatomical Connections Influence the Modulation of Conscious Visual Perception by High-Beta Frontal Oscillatory Activity. Cerebral cortex 25:2095–2101.

Quentin R, Chanes L, Migliaccio R, Valabregue R, Valero-Cabre A (2013) Fronto-tectal white matter connectivity mediates facilitatory effects of non-invasive neurostimulation on visual detection. Neuroimage 82:344–354.

Quentin R, Elkin Frankston S, Vernet M, Toba MN, Bartolomeo P, Chanes L, Valero-Cabre A (2016) Visual Contrast Sensitivity Improvement by Right Frontal High-Beta Activity Is Mediated by Contrast Gain Mechanisms and Influenced by Fronto-Parietal White Matter Microstructure. Cerebral cortex 26:2381–2390.

Romei V, Gross J, Thut G (2010) On the role of prestimulus alpha rhythms over occipito-parietal areas in visual input regulation: correlation or causation? J Neurosci 30:8692–8697.

Romei V, Thut G, Silvanto J (2016a) Information-Based Approaches of Noninvasive Transcranial Brain Stimulation. Trends Neurosci 39:782–795.

Romei V, Thut G, Mok RM, Schyns PG, Driver J (2012) Causal implication by rhythmic transcranial magnetic stimulation of alpha frequency in feature-based local vs. global attention. Eur J Neurosci 35:968–974.

Romei V, Bauer M, Brooks JL, Economides M, Penny W, Thut G, Driver J, Bestmann S (2016b) Causal evidence that intrinsic beta-frequency is relevant for enhanced signal propagation in the motor system as shown through rhythmic TMS. Neuroimage 126:120–130.

Rosanova M, Casali A, Bellina V, Resta F, Mariotti M, Massimini M (2009) Natural frequencies of human corticothalamic circuits. J Neurosci 29:7679–7685.

Rossi S, Hallett M, Rossini PM, Pascual-Leone A (2009) Safety, ethical considerations, and application guidelines for the use of transcranial magnetic stimulation in clinical practice and research. Clin Neurophysiol 120:2008–2039.

Ruff CC, Blankenburg F, Bjoertomt O, Bestmann S, Freeman E, Haynes JD, Rees G, Josephs O, Deichmann R, Driver J (2006) Concurrent TMS-fMRI and psychophysics reveal frontal influences on human retinotopic visual cortex. Curr Biol 16:1479–1488.

Ruzzoli M, Soto-Faraco S (2014) Alpha stimulation of the human parietal cortex attunes tactile perception to external space. Curr Biol 24:329–332.

Saalmann YB, Kastner S (2011) Cognitive and perceptual functions of the visual thalamus. Neuron 71:209–223.

Silvanto J, Lavie N, Walsh V (2006) Stimulation of the human frontal eye fields modulates sensitivity of extrastriate visual cortex. J Neurophysiol 96:941–945.

Stanislaw H, Todorov N (1999) Calculation of signal detection theory measures. Behav Res Methods Instrum Comput 31:137–149.

Steiger JH (1980) Tests for comparing elements of a correlation matrix. Psychological Bulletin 87:245–251.

Stewart LM, Walsh V, Rothwell JC (2001) Motor and phosphene thresholds: a transcranial magnetic stimulation correlation study. Neuropsychologia 39:415–419.

Stokes MG, Chambers CD, Gould IC, Henderson TR, Janko NE, Allen NB, Mattingley JB (2005) Simple metric for scaling motor threshold based on scalp-cortex distance: application to studies using transcranial magnetic stimulation. J Neurophysiol 94:4520–4527.

Suckling J, Bullmore E (2004) Permutation tests for factorially designed neuroimaging experiments. Hum Brain Mapp 22:193–205.

Tallon-Baudry C, Bertrand O (1999) Oscillatory gamma activity in humans and its role in object representation. Trends Cogn Sci 3:151–162.

Taylor PC, Nobre AC, Rushworth MF (2007) FEF TMS affects visual cortical activity. Cerebral cortex 17:391–399.

ter Braack EM, de Vos CC, van Putten MJ (2015) Masking the Auditory Evoked Potential in TMS-EEG: A Comparison of Various Methods. Brain Topogr 28:520–528.

Thut G, Pascual-Leone A (2010) Integrating TMS with EEG: How and what for? Brain Topogr 22:215–218.

Thut G, Schyns PG, Gross J (2011a) Entrainment of perceptually relevant brain oscillations by non-invasive rhythmic stimulation of the human brain. Front Psychol 2:170.

Thut G, Nietzel A, Brandt SA, Pascual-Leone A (2006) Alpha-band electroencephalographic activity over occipital cortex indexes visuospatial attention bias and predicts visual target detection. J Neurosci 26:9494–9502.

Thut G, Ives JR, Kampmann F, Pastor MA, Pascual-Leone A (2005) A new device and protocol for combining TMS and online recordings of EEG and evoked potentials. J Neurosci Methods 141:207–217.

Thut G, Veniero D, Romei V, Miniussi C, Schyns P, Gross J (2011b) Rhythmic TMS causes local entrainment of natural oscillatory signatures. Curr Biol 21:1176–1185.

Thut G, Bergmann TO, Frohlich F, Soekadar SR, Brittain JS, Valero-Cabre A, Sack AT, Miniussi C, Antal A, Siebner HR, Ziemann U, Herrmann CS (2017) Guiding transcranial brain stimulation by EEG/MEG to interact with ongoing brain activity and associated functions: A position paper. Clin Neurophysiol 128:843–857.

Van Opstal F, Van Laeken N, Verguts T, van Dijck JP, De Vos F, Goethals I, Fias W (2014) Correlation between individual differences in striatal dopamine and in visual consciousness. Curr Biol 24:R265–266.

Vernet M, Thut G (2014) Electroencephalography during Transcranial Magnetic Stimulation: current modus operandi. In: Neuromethods: Transcranial Magnetic Stimulation (Pascual-Leone A, Horvath JC, Rotenberg A, eds).

Vernet M, Brem AK, Farzan F, Pascual-Leone A (2014a) Synchronous and opposite roles of the parietal and prefrontal cortices in bistable perception: A double-coil TMS-EEG study. Cortex 64C:78–88.

Vernet M, Quentin R, Chanes L, Mitsumasu A, Valero-Cabre A (2014b) Frontal eye field, where art thou? Anatomy, function, and non-invasive manipulation of frontal regions involved in eye movements and associated cognitive operations. Frontiers in integrative neuroscience 8:66.

Vernet M, Bashir S, Yoo WK, Perez JM, Najib U, Pascual-Leone A (2013) Insights on the neural basis of motor plasticity induced by theta burst stimulation from TMS-EEG. Eur J Neurosci 37:598–606.

Wyart V, Tallon-Baudry C (2009) How ongoing fluctuations in human visual cortex predict perceptual awareness: baseline shift versus decision bias. J Neurosci 29:8715–8725.

Zaehle T, Rach S, Herrmann CS (2010) Transcranial alternating current stimulation enhances individual alpha activity in human EEG. PLoS One 5:e13766.

